# Cromolyn inhibits PGE_2_-mediated sensitisation of TRPV1 in a GPR35-dependent manner in sensory neurons

**DOI:** 10.64898/2026.01.21.700368

**Authors:** James P Higham, Luke W Paine, Alanna Cameron, Wendy Winchester, Ewan St John Smith, Naren Srinivasan, Rie Suzuki, James RF Hockley, David C Bulmer

## Abstract

There is a pressing need for effective alternatives to opioid analgesics, the development of which requires the identification of novel anti-nociceptive drug targets. Here, we have further investigated the anti-nociceptive properties of a GPR35 agonist, cromolyn, in an *in vitro* model of inflammatory sensitisation. We used ratiometric Ca^2+^ imaging of cultured sensory neurons to examine the effect of cromolyn on prostaglandin E_2_ (PGE_2_)-mediated sensitisation of the pro-nociceptive ion channel, transient receptor potential cation channel, subfamily V, member 1 (TRPV1). The sensitisation of TRPV1 by PGE_2_ was inhibited by cromolyn in a GPR35-dependent manner. These observations provide further evidence in support of an anti-nociceptive role for GPR35, highlighting the potential use of GPR35 agonists as analgesics.

## Introduction

The use of opioids as analgesics is associated with nausea, constipation, pruritus and sedation, and carries with it the risk of dependence with long-term use. The development of new analgesics requires a more detailed understanding of the mediators and mechanisms underpinning nociception, as well as the identification and validation of novel anti-nociceptive targets.

Inflammatory pain is characterised by hyper-sensitivity to noxious and innocuous stimuli (hyperalgesia and allodynia, respectively), and is mediated by the stimulation and sensitisation of nociceptors by inflammatory mediators (Kidd & Urban, 2001). Prostaglandin E_2_ (PGE_2_), a prototypical inflammatory mediator, sensitises nociceptors by potentiating the activity of ion channels central to nociceptor function, such as voltage-gated Na^+^ channels (England et al., 1996; Gold et al., 1996; Rush & Waxman, 2004), and the heat- and acid-sensitive cation channel, transient receptor potential cation channel, subfamily V, member 1 (TRPV1) (Lopshire & Nicol, 1998; Moriyama et al., 2005). The mechanism by which PGE_2_, acting through the EP_4_ receptor, sensitises TRPV1 is well-characterised. Sensitisation relies upon cyclic adenosine monophosphate (cAMP) activating protein kinase A which, in turn, phosphorylates TRPV1, orchestrated by AKAP79/150 (Schnizler et al., 2008; Zhang et al., 2008). This leads to an increase in TRPV1 surface expression (Ma et al., 2017) and possibly an increase in channel open probability (Lopshire & Nicol, 1998). TRPV1 is also sensitised by other inflammatory mediators, such as bradykinin and nerve growth factor (Chuang et al., 2001; Planells-Cases et al., 2005; Zhang et al., 2005), and plays a key role in thermal and mechanical hypersensitivity in models of inflammatory hyperalgesia (Flynn et al., 2014).

Transcriptomic data indicate that GPR35, TRPV1 and EP_4_ are co-expressed in murine and human sensory neurons (Bhuiyan et al., 2024). Given that GPR35 has been shown to couple to Gα_i/o_ G proteins (Gupta et al., 2025), leading to a reduction in cAMP signalling (Price et al., 2025), we hypothesised that GPR35 agonists may inhibit PGE_2_-mediated TRPV1 sensitisation. To test this hypothesis, we performed Ca^2+^ imaging of cultured murine sensory neurons. We found that PGE_2_-mediated TRPV1 sensitisation was inhibited by pre-treatment with the GPR35 agonist, cromolyn. We also tested whether this effect of cromolyn required GPR35. Indeed, in sensory neurons lacking GPR35, cromolyn failed to inhibit PGE_2_-mediated TRPV1 sensitisation. These observations provide further support for an anti-nociceptive role for GPR35 in sensory neurons.

## Methods

### Preparation of sensory neurons

Sensory neurons were isolated from dorsal root ganglia (DRG) harvested from male mice culled between 8-16 weeks of age. C57Bl/6 mice were obtained from Charles River (RRID: ISMR_JAX:000664) and GPR35^+/+^ and GPR35^-/-^ mice (C57BL/6-Gpr35tm1b(EUCOMM)Hmgu/WtsiH) were rederived by MRC Harwell and INFRAFRONTIER/EMMA. Mice were housed in groups of up to five littermates with *ad libitum* access to food, water, bedding and enrichment (e.g., shelters, tunnels, chews). All animal work was ethically reviewed and carried out in accordance with Schedule 1 of the Animals (Scientific Procedures) Act 1986 Amendment Regulations 2012 and the GSK Policy on the Care, Welfare and Treatment of Laboratory Animals.

After harvesting, DRG were enzymatically dissociated by incubation with Lebovitz L-15 GlutaMAX medium (Invitrogen, supplemented with 24 mM NaHCO_3_ and 6 mg/mL bovine serum albumin) containing 1 mg/mL collagenase (Sigma Aldrich) for 15 minutes, followed by 1 mg/mL trypsin (Sigma Aldrich) for 30 minutes (at 37°C/5% CO_2_). Following this, DRG were transferred to Lebovitz L-15 GlutaMAX culture medium supplemented with 10% foetal bovine serum (Gibco), 24 mM NaHCO_3_ (Gibco), 38 mM glucose (Gibco) and 2% vol/vol penicillin/streptomycin (Gibco). DRG were mechanically dissociated through multiple rounds of gentle trituration through a P1000 pipette tip. After dissociation, dissociated neurons were pelleted, resuspended in the above culture medium and seeded on to 35 mm glass-bottomed dishes (MatTek) coated with poly-D-lysine and laminin (Gibco). Neurons were incubated at 37°C/5% CO_2_ until use.

### Ca^2+^ imaging of sensory neurons

Neurons were used for experiments 16-24 hours after seeding. After washing to remove culture medium, neurons were loaded with Fura2 by incubation with 2 μM Fura2-AM (Invitrogen) for 45 minutes at room temperature (shielded from light). Neurons were washed again, bathed in extracellular buffer (containing, in mM: 140 NaCl, 4 KCl, 1 MgCl_2_, 2 CaCl_2_, 4 D-glucose, and 10 HEPES; pH adjusted to 7.4 using NaOH) and visualised with a 20x objective (Olympus UApo/340) on an Olympus IX71 inverted microscope. Neurons were continuously superfused with extracellular buffer using a flexible perfusion pencil (Automate Scientific) fed by a valve-controlled perfusion system (Biologic RSC-200). Following a 10 second baseline, capsaicin (100 nM) was applied for 20 seconds, after which either vehicle (0.01% DMSO), PGE_2_ (1 μM) or PGE_2_ and cromolyn (10 μM) were applied for 3 minutes. Following treatment, capsaicin was applied again and, finally, KCl (50 mM) was applied for 10 seconds to identify viable neurons.

Fura2 was alternately excited at 340 nm and 380 nm using LED light sources (Cairn Research), and emission at 520 nm was captured at 1 fps with 100 ms exposure using a Hamamatsu OrcaFlash 4.0LT CCD camera. Data were recorded and initially analysed using MetaFluor (Molecular Devices). Regions of interest were manually traced around neurons and the average intensity of emission at 520 nm following excitation at 340 nm and 380 nm was calculated. The ratio of these emission intensities (F_340_/F_380_, referred to as “R”) provides a metric of intracellular Ca^2+^ concentration. F_340_/F_380_ was normalised to its baseline value (R/R_0_). Only neurons which exhibited a stable baseline and an increase in F_340_/F_380_ of >10% over baseline during KCl application were included in analysis. Capsaicin-sensitive neurons were identified using the same cutoff. The magnitude of capsaicin-evoked Ca^2+^ transients was found by subtracting baseline R/R_0_ (immediately prior to capsaicin application) from peak R/R_0_ (see Figure 1A which shows that the baseline may change between the first and second application of capsaicin).

**Figure 1:**
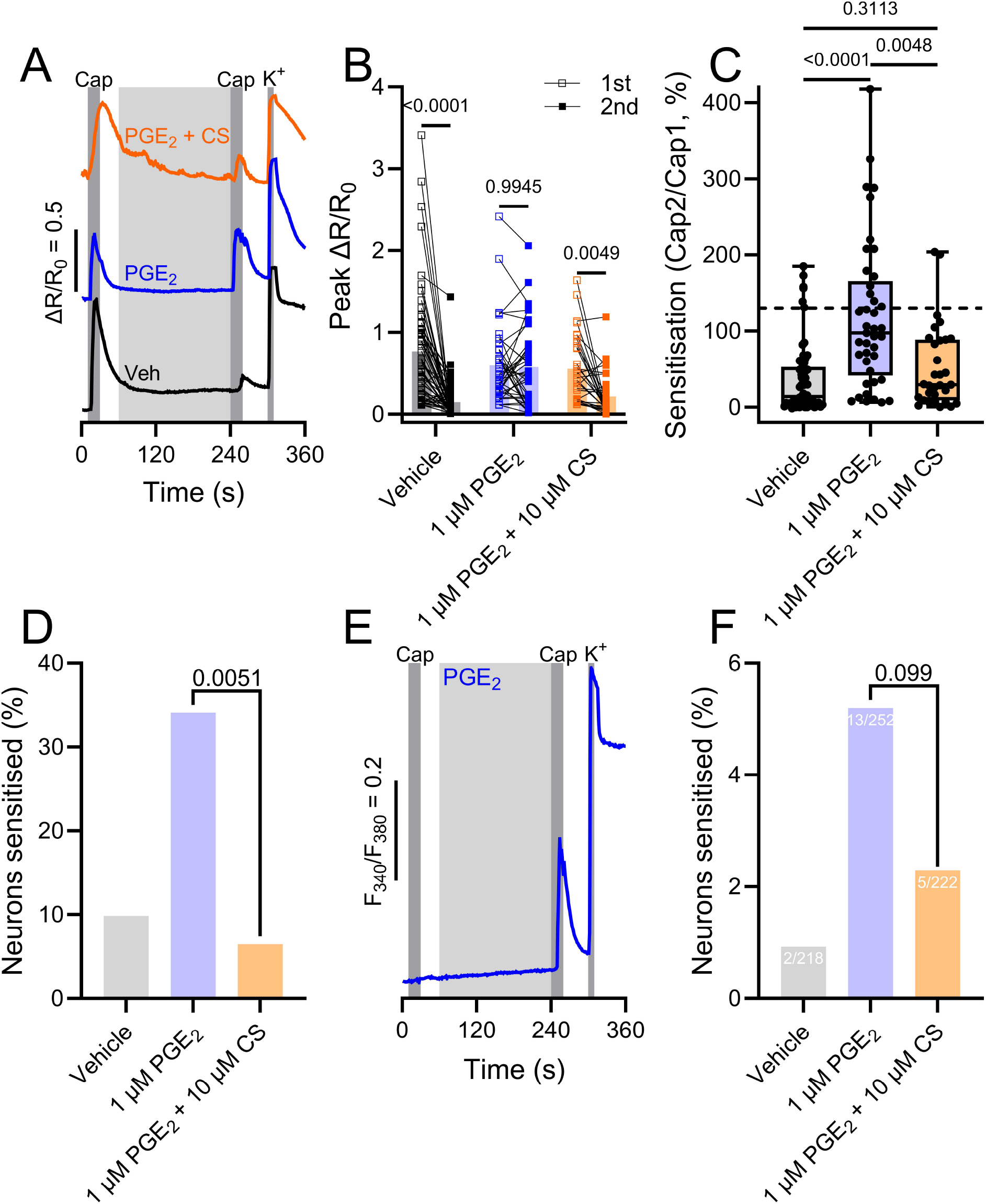
Inhibition of PGE_2_-mediated TRPV1 sensitisation by cromolyn. (A) Example traces showing the experimental protocol and the effect of vehicle (black), PGE_2_ (blue) or PGE_2_ and cromolyn (orange) on the tachyphylaxis of the response to capsaicin. Notice the marked tachyphylaxis of the response to capsaicin in the vehicle-treated neuron, which is absent in the PGE_2_-treated neuron. (B) Grouped data showing the magnitude of the response to the first (open squares) and second (filled squares) applications of capsaicin under different conditions (3 min incubation with vehicle, PGE_2_ or PGE_2_ and cromolyn). Bars show the mean for each group. Data analysed using a repeated measures two-way ANOVA with Sidak’s post-hoc tests. (C) Grouped data showing the ratio of the magnitude of the second response to capsaicin to that of the first (Cap2/Cap1) under different conditions, as a metric of sensitisation. Plots show the mean, interquartile range and full range for each group. Dashed line shows the mean plus two standard deviations of the vehicle-treated group. Data analysed using a Kruskal-Wallis test with Dunn’s post-hoc tests. (D) The fraction of capsaicin-sensitive neurons (in C) in which the response to capsaicin was sensitised under different conditions. Data analysed using Chi-squared tests. (E) Example trace showing a neuron which was initially insensitive to capsaicin, but became sensitive following the application of PGE_2_. (F) The fraction of neurons initially insensitive to capsaicin, but which became sensitive following the application of vehicle, PGE_2_ or PGE_2_ and cromolyn. Data analysed using Chi-squared tests.

### Statistics

Data were scrutinized to ensure they met the assumptions of parametric analyses. Normality was assessed using the Shapiro-Wilk test and homogeneity of variances using the F-test. Where appropriate, non-parametric, rank-based statistical tests were used. Data are presented as mean ± standard error or median with interquartile range (IQR). Details of statistical tests used are provided in the figure legends, and sample sizes (number of neurons) are provided in the text. In all experiments, neurons were derived from 3-4 independent biological replicates. Data were plotted and analysed using GraphPad Prism (GraphPad Software, v10.2.0).

## Results

### Cromolyn inhibits PGE_2_-mediated sensitisation of TRPV1

We examined the effect of the inflammatory mediator, PGE_2_, on TRPV1 channel function. To do this, we applied capsaicin (a TRPV1 agonist; 100 nM, 20 s) to Fura2-loaded sensory neurons (from wild-type C57Bl/6 mice) twice and, in the time between applications, treated neurons for three minutes with either vehicle (0.01% DMSO), PGE_2_ (1 μM) or PGE_2_ and cromolyn (10 μM) (Figure 1A). A three minute application was chosen as it allows marked tachyphylaxis of the response to capsaicin to be observed (Schnizler et al., 2008). In control conditions, the response to capsaicin underwent significant tachyphylaxis (Figure 1A), with the response to the second application significantly reduced compared to the first (n = 61, p < 0.0001, Figure 1B). Treatment with PGE_2_ abolished tachyphylaxis, and the response to the second application of capsaicin was indistinguishable from the first (n = 41, p = 0.99, Figure 1A and B). Treatment with both PGE_2_ and cromolyn prevented the effect of PGE_2_, with the response to the second application of capsaicin being significantly reduced compared to the first (n = 31, p = 0.0049, Figure 1A and B).

The response to the first application of capsaicin was, as expected, no different between groups (p > 0.22), indicating that the effect of the respective treatments was solely on the response to the second application of capsaicin. Indeed, the response to the second application of capsaicin was larger in PGE_2_-treated neurons compared to vehicle-treated neurons (p < 0.0001), while this response was reduced in neurons treated with PGE_2_ and cromolyn compared to those treated with PGE_2_ alone (p = 0.0094). There was no difference in the magnitude of the second response to capsaicin in neurons treated with vehicle and those treated with PGE_2_ and cromolyn (p = 0.99).

To further quantify the effect of PGE_2_ and cromolyn, the ratio of the response to the second and first application of capsaicin was calculated (Cap2/Cap1). In vehicle-treated neurons, this ratio was 14.0% (median; IQR, 3.9-52.9%), indicating substantial tachyphylaxis (Figure 1C). In PGE_2_-treated neurons, the response ratio was 97.6% (IQR, 41.6-165.6%, p < 0.0001 compared to vehicle-treated neurons), showing an absence of tachyphylaxis due to an increase in the second response (Figure 1C). In neurons treated with PGE_2_ and cromolyn, the response ratio was 29.3% (IQR, 8.7-88.8%), indicating an inhibition of the effect of PGE_2_ (p = 0.0048 compared to PGE_2_ alone, p = 0.31 compared to vehicle, Figure 1C). The proportion of capsaicin-sensitive neurons in which the response to the second application of capsaicin was sensitised (Cap2/Cap1 greater than the mean plus two standard deviations of the control group) was 34.1% in PGE_2_-treated neurons, compared to 9.8% (p = 0.0024) and 6.5% (p = 0.0051) in neurons treated with vehicle or PGE_2_ and cromolyn, respectively (Figure 1D).

In a very small fraction of all neurons (2/218), a response to the second application of capsaicin was observed in the absence of any response to the first application. The fraction of neurons exhibiting this pattern of responses was increased by treatment with PGE_2_ (13/252, p = 0.0091, Figure 1E and F). The fraction of neurons exhibiting this pattern of responses in neurons treated with PGE_2_ and cromolyn was no different to those treated with vehicle (5/222, p = 0.26) nor those treated with PGE_2_ alone (p = 0.099, Figure 1F).

### The effect of cromolyn is dependent on GPR35

Next, we examined the effect of cromolyn on PGE_2_-mediated sensitisation of TRPV1 in neurons derived from GPR35 knock-out (GPR35^-/-^) mice and their GPR35^+/+^ littermates. Figure 2A shows example traces from GPR35^+/+^ neurons, which behaved similarly to neurons derived from wild-type C57Bl/6 mice. In vehicle-treated neurons, marked tachyphylaxis of the response to capsaicin was observed (n = 38, p < 0.0001, Figure 2B). Treatment with PGE_2_ abolished tachyphylaxis of the response to capsaicin, with the first and second responses being of similar magnitude (n = 37, p = 0.15, Figure 2B). This effect of PGE_2_ was inhibited by cromolyn and a marked reduction in the response to capsaicin between the first and second application was observed (n = 46, p < 0.0001, Figure 2B). The magnitude of the response to the second application of capsaicin was larger in PGE_2_-treated neurons compared to vehicle-treated neurons (p < 0.0001). Treatment with PGE_2_ and cromolyn reduced the magnitude of the second response (p < 0.0001 compared to PGE_2_ treatment alone, p = 0.50 compared to vehicle treatment).

**Figure 2:**
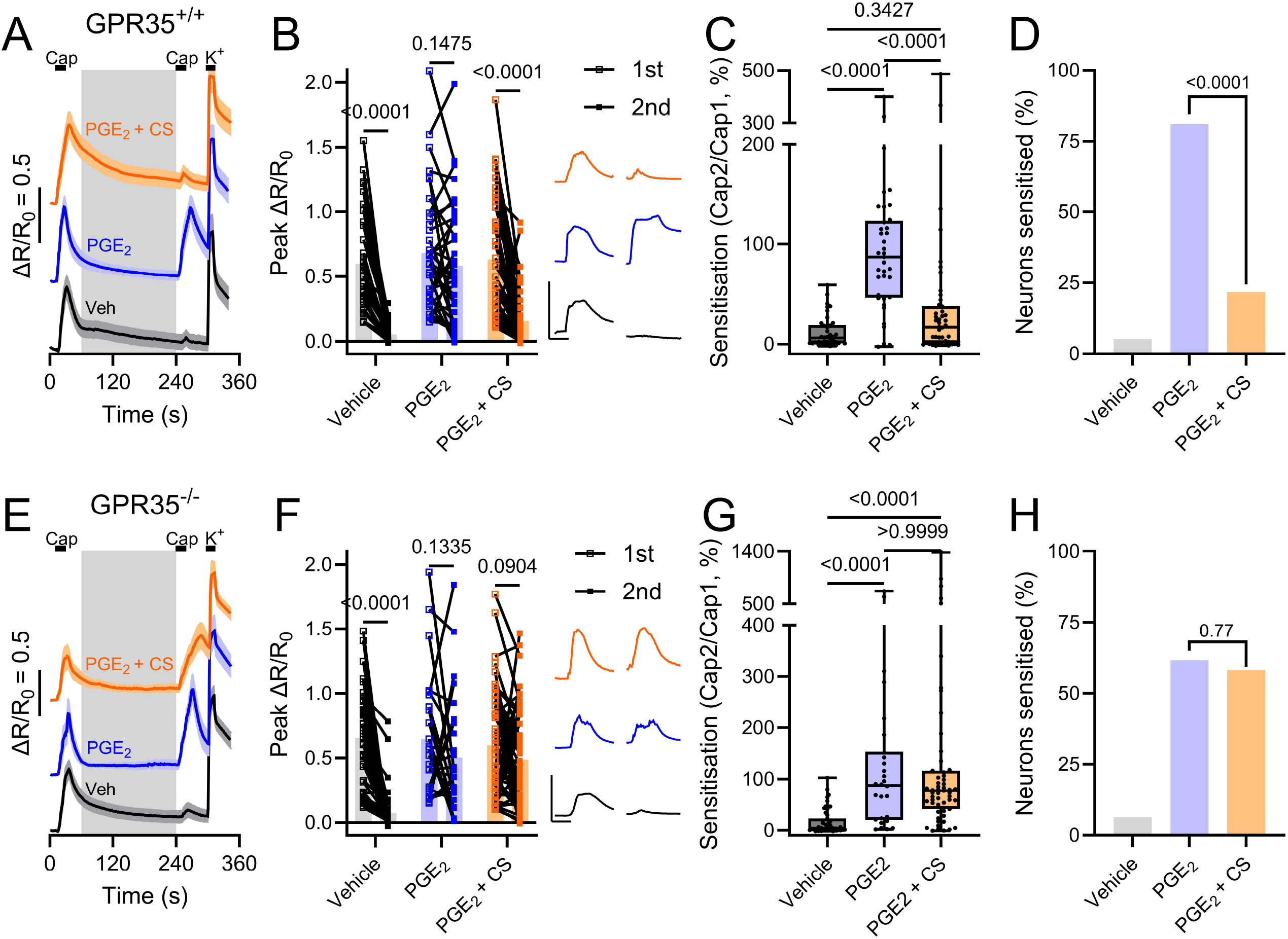
The inhibitory effect of cromolyn is dependent on GPR35. (A) Example traces showing the effect of vehicle (black), PGE_2_ (blue) or PGE_2_ and cromolyn (orange) on the tachyphylaxis of the response to capsaicin in GPR35^+/+^ neurons. Solid line is the mean and the shaded area is the standard error; neurons in each group taken from the same experiment. (B) Grouped data showing the magnitude of the response to the first (open squares) and second (filled squares) applications of capsaicin under different conditions (3 min incubation with vehicle, PGE_2_ or PGE_2_ and cromolyn) in GPR35^+/+^ neurons. Bars show the mean for each group. Data analysed using a repeated measures two-way ANOVA with Sidak’s post-hoc tests. Inset traces show raw traces of F_340_/F_380_ values for the first (left) and second (right) applications of capsaicin under different conditions. Scale: 0.2 F_340_/F_380_; 20 s. (C) Grouped data showing the ratio of the magnitude of the second response to capsaicin to that of the first (Cap2/Cap1) under different conditions in GPR35^+/+^ neurons. Plots show the mean, interquartile range and full range for each group. Data analysed using a Kruskal-Wallis test with Dunn’s post-hoc tests. (D) The fraction of capsaicin-sensitive neurons (in C) in which the response to capsaicin was sensitised under different conditions in GPR35^+/+^ neurons. Data analysed using Chi-squared tests. (E) Example traces showing the effect of vehicle (black), PGE_2_ (blue) or PGE_2_ and cromolyn (orange) on the tachyphylaxis of the response to capsaicin in GPR35^-/-^ neurons. Solid line is the mean and the shaded area is the standard error; neurons in each group taken from the same experiment. (F) Grouped data showing the magnitude of the response to the first (open squares) and second (filled squares) applications of capsaicin under different conditions (3 min incubation with vehicle, PGE_2_ or PGE_2_ and cromolyn) in GPR35^-/-^ neurons. Bars show the mean for each group. Data analysed using a repeated measures two-way ANOVA with Sidak’s post-hoc tests. Inset traces show raw traces of F_340_/F_380_ values for the first (left) and second (right) applications of capsaicin under different conditions. Scale: 0.2 F_340_/F_380_; 20 s. (G) Grouped data showing the ratio of the magnitude of the second response to capsaicin to that of the first (Cap2/Cap1) under different conditions in GPR35^-/-^ neurons. Plots show the mean, interquartile range and full range for each group. Data analysed using a Kruskal-Wallis test with Dunn’s post-hoc tests. (H) The fraction of capsaicin-sensitive neurons (in G) in which the response to capsaicin was sensitised under different conditions in GPR35^-/-^ neurons. Data analysed using Chi-squared tests.

In vehicle-treated neurons, Cap2/Cap1 was 6.3% (IQR, 0.9-19.0%), compared to 87.1% (IQR, 46.2-123.0%; p < 0.0001) and 16.9% (IQR, 0.9-37.8%; p = 0.34) in neurons treated with PGE_2_ alone and PGE_2_ and cromolyn, respectively (Figure 2C). Cap2/Cap1 was reduced by treatment with PGE_2_ and cromolyn compared to treatment with PGE_2_ alone (p < 0.0001, Figure 2C). The fraction of neurons in which the response to the second application of capsaicin was sensitised was 81.1% in neurons treated with PGE_2_, compared to 21.7% in neurons treated with PGE_2_ and cromolyn (p < 0.0001, Figure 2D).

In contrast to neurons isolated from wild-type and GPR35^+/+^ mice, in GPR35^-/-^ neurons, cromolyn failed to inhibit PGE_2_-mediated sensitisation of TRPV1 (Figure 2E). The tachyphylaxis of the response to capsaicin, and PGE_2_-induced sensitisation, was similar between GPR35 WT and KO neurons. In control conditions, marked tachyphylaxis of the response to capsaicin was observed in GPR35^-/-^ neurons (n = 47, p < 0.0001, Figure 2F). Tachyphylaxis was absent following treatment with PGE_2_ (n = 26, p = 0.13, Figure 2F). In contrast to GPR35^+/+^ neurons, there was no effect of cromolyn in GPR35^-/-^ neurons – the response to the second application of capsaicin remained sensitised, and there was no difference in the magnitude of the first and second response (n = 55, p = 0.090, Figure 2F). Treatment with PGE_2_ increased the magnitude of the second response to capsaicin compared to vehicle treatment (p < 0.0001). In neurons treated with PGE_2_ and cromolyn, the magnitude of the second response was no different compared to neurons treated with PGE_2_ alone (p > 0.99) and remained greater than in neurons treated with vehicle (p < 0.0001).

Cap2/Cap1 was increased from 2.9% (IQR, 1.0-23.1%) in vehicle-treated neurons to 87.8% (IQR, 20.1-153.1%) in PGE_2_-treated neurons (p < 0.0001, Figure 2G). In neurons treated with PGE_2_ and cromolyn, Cap2/Cap1 was 76.8% (IQR, 41.1-116.5%), no different to PGE_2_ treatment alone (p > 0.99), and greater than vehicle treatment (p < 0.0001, Figure 2G). The fraction of neurons in which the response to the second application of capsaicin was sensitised was 61.5% in neurons treated with PGE_2_, compared to 58.2% in neurons treated with PGE_2_ and cromolyn (p = 0.77, Figure 2H).

## Discussion

This study has shown that cromolyn blocked TRPV1 sensitisation induced by the pro-nociceptive inflammatory mediator, PGE_2_. Others have shown morphine to have a similar effect on TRPV1 sensitisation (Vetter et al., 2006, 2008). Experiments using sensory neurons lacking GPR35 indicated that cromolyn acted through this receptor. This extends our previous work wherein we showed that the stimulation of GPR35 inhibited colonic afferent hypersensitivity to mechanical stimuli (Gupta et al., 2025).

The data presented here are in agreement with a previous study showing that kynurenic acid and zaprinast, an endogenous and a synthetic GPR35 agonist, respectively, inhibited PGE_2_-induced sensory neuron hyperexcitability (Resta et al., 2016). It was proposed that this effect was due to inhibition of the depolarising shift in the voltage-dependence of activation of HCN channels caused by PGE_2_-mediated cAMP production (Resta et al., 2016). Both GPR35 agonists also ameliorated PGE_2_-induced inflammatory hyperalgesia, demonstrating the analgesic effects of GPR35 stimulation (Cosi et al., 2011; Resta et al., 2016). It has also been reported that kynurenic acid and zaprinast inhibited voltage-gated Ca^2+^ channel current in sympathetic neurons which heterologously expressed GPR35 (Guo et al., 2008). Ca^2+^ current inhibition was blocked by a depolarising pre-pulse, indicating that this effect is likely mediated by βγ G-protein subunits, as is the case with other Gα_i_-coupled receptors (Raingo et al., 2007; Seward et al., 1991). These observations indicate that GPR35 signals through multiple anti-nociceptive pathways, leading to a suppression of nociceptive signalling.

We found that treatment with PGE_2_ unmasked a small “silent” capsaicin-sensitive population of sensory neurons. Other inflammatory insults have been shown to increase the fraction of sensory neurons which express TRPV1. For example, the induction of inflammation in the knee using Freud’s Complete Adjuvant increased the fraction of knee-innervating neurons sensitive to capsaicin and the fraction expressing TRPV1 (Chakrabarti et al., 2018). Moreover, an elevated fraction of TRPV1-expressing sensory neurons has been found following partial sciatic nerve ligation (PSNL) – a model of neuropathic pain (Ma et al., 2010). Levels of PGE_2_ were also increased in the PSNL model. The administration of a cyclooxygenase 2 inhibitor following PSNL attenuated the elevation in the expression of TRPV1 and the amount of PGE_2_ (Ma et al., 2010). Perhaps, therefore, PGE_2_ contributes to an increase in TRPV1 expression and/or function following inflammatory or neuropathic insults. What’s more, in colonic biopsies from human patients with irritable bowel syndrome, PGE_2_ was shown to be increased in abundance (Cenac et al., 2015; Grabauskas et al., 2020) and so too was the density of TRPV1-expressing sensory nerve fibres (Akbar et al., 2008).

This study provides further evidence in support of an anti-nociceptive role for GPR35. Future investigations should focus on further elucidating the analgesic properties of GPR35 agonists in models of inflammatory pain and, more importantly, in human inflammatory diseases.

